# *Wolbachia* facilitates the reproduction of a parthenogenetic ladybug

**DOI:** 10.64898/2026.06.16.732365

**Authors:** Kristine Jecha, Nathalie Parthuisot, Emilie Lecompte, Tanja Schwander, Alexandra Magro

**Author notes:** Co-last authors (Tanja Schwander, Alexandra Magro).

## Abstract

*Wolbachia* is a bacterial endosymbiont that is primarily transmitted from mother to offspring. To increase their transmission, some strains manipulate their host’s reproduction to favor female offspring, such as by inducing parthenogenesis. Here, we assess whether *Wolbachia* induces parthenogenesis in recently discovered parthenogenetic populations of the ladybug *Nephus voeltzkowi* by treating females with an antibiotic. Females from sexual populations, which we show are not infected by *Wolbachia*, serve as a control. Our results demonstrate that the treatment decreases *Wolbachia* load, subsequently reducing egg production and development among parthenogenetic females, while having no effect on sexual females. *Wolbachia* load and reproduction then rebound when the treatment is removed. This suggests that the *Wolbachia* infection is necessary for successful reproduction in the parthenogenetic females, and it may play a two-part role by facilitating egg laying and late embryo development. The cooccurrence of *Wolbachia* infection and parthenogenesis in *N. voeltzkowi*, as well as *Wolbachia*’s manipulation of host reproduction makes this a likely candidate of *Wolbachia*-induced parthenogenesis outside of haplo-diploids. Understanding how *Wolbachia* can impact diverse insect reproductive systems can shed a light on the extent in which these bacteria can manipulate hosts for their own gain.

## Introduction

*Wolbachia* is a bacterial endosymbiont that is widespread across Arthropoda, with estimations of 50% to76% of arthropod species being infected (Jeyaprakash & Hoy, 2000; Weinert et al., 2015). *Wolbachia* bacteria live within the cytoplasm of the host cells, and their primary method of transmission is via the cytoplasm of the egg from the mother. Due to this transmission strategy, male hosts are a dead end and some *Wolbachia* strains have instituted various mechanisms to manipulate their host’s reproductive process to increase transmission (Werren et al., 1995, 2008). Such manipulation includes *Wolbachia*-induced male-killing, cytoplasmic incompatibility, feminization, and thelytokous parthenogenesis (Barr, 1980; Hurst, Jiggins, Schulenburg, et al., 1999; Rousset et al., 1997; Stouthamer & Werren, 1993). Thelytokous parthenogenesis (hereafter, referred to as “parthenogenesis”) is the production of female offspring by a mother without the contribution of a male. In contrast to other forms of reproductive manipulation, parthenogenesis induction is not expected to generate conflict with the host, as it does not reduce host fertility and instead confers a demographic advantage to infected females through the classical two-fold benefit of asexual reproduction.

*Wolbachia-*induced parthenogenesis has been very well documented in haplo-diploid insects (reviewed in Ma & Schwander, 2017). Haplo-diploids employ a form of sex determination wherein fertilized, diploid eggs develop as a female whereas unfertilized, haploid eggs develop as male. *Wolbachia* can interfere with the reproductive process by diploidizing these unfertilized eggs and, in some cases, feminizing them in a second step, generating female offspring without male contribution (Ma et al., 2015; Stouthamer & Kazmer, 1994; Verhulst et al., 2023). While it has been experimentally demonstrated in haplo-diploids, *Wolbachia* induced parthenogenesis has thus far not been confirmed in organisms with other sex determination systems. Evidence that *Wolbachia* is inducing parthenogenesis in its host can be found with the use of antibiotic or heat treatments to reduce or eliminate the *Wolbachia* infection (Stouthamer et al., 1990). In the case of insects with haplo-diploid sex determination, following treatments, the hosts can be capable of resuming sexual reproduction and the production of male offspring from unfertilized eggs. As other sex determination systems lack the clear evidence of the reversal of parthenogenesis induction, it is difficult to formally demonstrate *Wolbachia* induced parthenogenesis in diplo-diploids (Ma & Schwander, 2017).

While *Wolbachia* induced parthenogenesis has yet to be definitively proven in non-haplo-diploids, there are still many diplo-diploid parthenogenetic lineages that are hypothesized to be *Wolbachia* induced (reviewed in Ma & Schwander, 2017). In most of these cases, *Wolbachia* induced parthenogenesis is hypothesized because *Wolbachia* infection is detected in parthenogenetic but not sexual species or populations, but experimental assessments are scarce (in springtails: Frati et al., 2004; in weevils: da Cruz Cabral et al., 2024; Rodriguero et al., 2010). Further research on non-haplo-diploid, parthenogenetic species with *Wolbachia* infections is needed to understand whether and how parthenogenesis can be induced in different sex determination mechanisms. This opportunity presented itself by the discovery of parthenogenetic, *Wolbachia*-infected populations of the ladybug (Coccinellidae: Coleoptera) *Nephus voeltzkowi* Weise a species previously documented as sexually reproducing with bisexual populations (Magro et al., 2020). This finding may indicate a case of *Wolbachia*-induced parthenogenesis outside of haplo-diploids. Here we determine the status of *Wolbachia* infection in sexually reproducing individuals and investigate the role that *Wolbachia* is having on the reproduction of parthenogenetic females by administering an antibiotic to reduce the infection.

## Materials and Methods

### Stock cultures

Sexual and parthenogenetic stock cultures of *N. voeltzkowi* were initiated with adults collected in the field in 2023, with 30 parthenogenetic females collected in La Réunion island and 60 sexual individuals in the Sète region of southern France. Stock cultures were reared and maintained in 175 cm^3^ plastic boxes containing a piece of corrugated paper at 23 ± 1 °C, ∼78% humidity, and a LD 16:8 photoperiod, as previously described (Magro et al., 2020). All experiments were conducted with the same temperature, humidity, and photoperiod as the stock cultures. Cultures were fed *ad libitum* with coccids (*Planococcus citri* Risso) by adding coccid-infested, organic potato sprouts once a week to their boxes. To limit overpopulation, adults were collected each week and kept in batches of about 20 individuals in new 175 cm^3^ plastic boxes and fed as described above.

### *Wolbachia* elimination via antibiotic treatment

We used 163 adult females for assessing the role of *Wolbachia* in inducing parthenogenesis: 51 sexual females and 112 parthenogenetic females. They were split into two treatment groups, antibiotic diet and control diet. Parthenogenetic females in each treatment were further split into those paired with a male and those without, while all sexual females were paired with a male, see Table 1. For logistic reasons, we conducted the experiment in five separate batches, with approximately balanced sample sizes across the treatments in each batch, see Supplementary table S1.

**Table 1.** Sample size of *N. voeltzkowi* females from the different reproductive modes and treatment groups. See Supplementary table S1 for assignment of samples to batches.

|  | *Antibiotic diet* |  | *Control diet* |  |
| --- | --- | --- | --- | --- |
|  | Parthenogenetic | Sexual | Parthenogenetic | Sexual |
| *Without male* | 30 | - | 30 | - |
| *With male* | 26 | 24 | 26 | 27 |
| **Total** | **56** | **24** | **56** | **27** |

Females were isolated in *Drosophila* vials, ø x h = 25 x 95 mm closed with cellulose acetate vial plugs. Females were cyclically rotated every two days between vials containing an artificial diet (consisting of agar, vitamins, sugar, yeast, and dehydrated liver) adapted from Majerus & Kearns, 1989 and vials containing a coccid diet (consisting of a cutting of coccid-infested, organic potato sprout). This transfer is necessary because the use of artificial diets to support ladybug cultures has proved challenging (Hodek & Evans, 2012) and, to our knowledge, *Nephus* species cannot survive and reproduce exclusively on artificial diets. In addition to aiding *N. voeltzkowi* rearing, the period with coccid diet allows females from the relevant treatment groups access to a male with which to copulate so that the fecundity measurements obtained after being isolated in the rearing vials with artificial diet would not be affected by the presence of males, as well as to ensure males are not exposed to the antibiotic treatment. The antibiotic treatment was administered by adding Rifampicin antibiotic to the artificial diet, at a dose of 1 mg/ml. The same artificial diet excluding Rifampicin was used for the control treatment. A fresh ∼0.5 cm x 0.5 cm piece of treatment-appropriate artificial diet was added to their vials at the beginning of every artificial diet two-day period. At the end of this period, any eggs laid into the vial plug were counted under a Nikon SMZ1000 stereo microscope and incubated in the plug (see details below) and the females were transferred into a different set of vials containing a cutting of coccid-infested, organic potato sprout for two days. Vials were used for two cycles before being replaced with new vials to maintain cleanliness. Note that females usually lay eggs into coccid ovisacs (when available) but lay all eggs into the cellulose acetate vial plug in the absence of coccid material. The total treatment period was conducted over 32 days (corresponding to 8 two-day periods of artificial diet and 8 two-day periods of coccid diet).

Due to the limited number of males available, two females from the same treatment group were paired with the one male in a coccid-diet vial at the same time. In order to easily identify the 2 females and the male present in the coccid-diet vials, each individual was marked on its elytra with a different color of water-based Posca paint markers PC1-MR (Mitsubishi Pencil Co.). After 2 days, the females were cycled to their individual, artificial-diet vials and the males remained in their individual coccid vials during the artificial diet period until the females are cycled back in. In the event that a male died before the end of the experiment, a replacement male was added to the vial. If a replacement male was not available, the females would be removed from the experiment.

After the 32-day treatment period, half of the females from each treatment group (75 females, see Supplementary Table 1) were individually frozen at -21 °C to later quantify their *Wolbachia* titer (see below). The remaining females were either dissected to visually inspect their ovaries for presence or absence of oogenesis (18 females, see Supplementary Table 1) or entered the “observation period” where they were maintained similarly as described above with the exception that all treatment groups received only artificial diet without antibiotic, including those initially treated with the antibiotic (55 females, see Supplementary Table 1).

Throughout the experiment, any eggs laid by females that died or were removed before the end of the experiment (22 females, see Supplementary Table 1) were still counted and incubated as described below. At the end of the observation period (70 days after the start of the experiment), all remaining individuals were frozen at -21 °C for *Wolbachia* titer analysis, see below (48 females, see Supplementary Table 1).

### Egg incubation and collection

The collected plugs with eggs from each artificial diet period were placed in empty *Drosophila* vials and incubated for 4 days in the same conditions as described above, after which a small cutting of coccid-infested potato sprout was added to each vial to feed the soon-to-hatch larvae. Incubation then continued for 14 more days, after which we recorded the number of eggs that hatched, showed signs of development without hatching (larvae visible through the egg chorion), or showed no signs of development.

### Dissection and tissue staining

To visually inspect their reproductive system for potential effects of antibiotic treatment or *Wolbachia* elimination, we dissected 18 females across each treatment group one to three days after their eighth artificial diet period. Their reproductive system was stained by adding an aqueous cresyl blue solution (100ml distilled water, 1g Brilliant cresyl blue powder, 0.5g Sodium Dodecyl Sulfate) to the tissue for 5-10 minutes before removing the solution and gently rinsing the tissue with distilled water. This allowed us to clearly examine the reproductive system under a Nikon SMZ1000 stereo microscope for evidence of egg production at the end of the treatment period of the experiment. Signs of egg production included the presence of distinct eggs in the ovary or vaginal canal or presence of swollen ovariole follicles.

### Incomplete *Wolbachia* treatment

To develop more detailed insights into how *Wolbachia* influences reproduction at different titer levels, we initiated a second, simplified experiment with 16 parthenogenetic females. The experiment followed the same diet transfer routine as the main experiment described above for unmated parthenogenetic females for only two cycles of antibiotic treatment. 10 females were given an artificial diet with antibiotic, and 6 females were supplied an artificial diet without antibiotic. After the second artificial diet cycle (6 days), 8 eggs from 5 treated females and 5 eggs from 1 control female were collected from the vial plugs to verify early embryo development via DAPI staining and females were frozen at -21 °C for *Wolbachia* titer quantification (described below).

To prepare the eggs for DAPI staining, eggs laid into the vial plugs during their last 2-day artificial diet period were removed using a small paint brush and placed on black membrane polycarbonate 0.22 µm filter paper which was place on a GFF 0.7 μm filter on a vacuum flask. A glass funnel was affixed on top of the filters, and the eggs were stained by directly applying 25 drops of a 0.01 mg m/L DAPI stain and 5mL of Milli-Q water for 15min before removing the liquid with suction. The filter paper was then transferred to a microscope slide, and the eggs were flattened using a glass microscope slide cover. These were then observed to determine any evidence of egg development by the presence of fluorescent nuclei using a Nikon Eclipse 80 microscope, equipped with a Mercure Nikon C-LHGFI HG lamp and a DAPI filter (excitation filter: 362-396 nm, barrier filter: 432-482 nm, dichroic mirror: 415 nm).

### *Wolbachia* titer quantification

The relative levels of *Wolbachia* titer in the subset of parthenogenetic females (from three time points: after the second artificial diet cycle, after the treatment period, and after the observation period) and sexual females (after treatment period and after observation period) were quantified using digital PCR (dPCR) based on the ratio of the single-copy *Wolbachia* surface protein (*wsp*) gene to the single-copy *N. voeltzkowi* actin gene.

The elytra and head of each female were clipped off, and DNA extractions were carried out on the remaining body material using the QIAamp DNA Micro Kit (Qiagen, Germany). Each individual was placed in 180 µL ATL buffer with 20 µL Proteinase K, then crushed using a sterile piston in a 1.5 mL microtube. Samples were incubated at 56 °C overnight. The remainder of the protocol followed the manufacturer’s instructions. DNA was finally eluted in 55 µL AE buffer.

We developed two dPCR oligo sets, each with two primers and a fluorescently labeled probe (Table 2). We targeted the single-copy *Wolbachia wsp* gene with the single-copy *N. voeltzkowi* actin gene as our endogenous control gene. Oligo sequences, based on the actin gene sequence of *N. voeltzkowi* and *wsp* gene of its *Wolbachia* strain, were obtained by mapping the paired-end reads short reads produced previously (Magro et al., 2020) onto reference sequences available in a public database (accession numbers MK097204 and AJ130714 for actin and *wsp* gene, respectively) using Geneious v.9.1.7 (https://www.geneious.com) and the default parameters. Assemblages were verified and consensus sequences were extracted. We then generated new assemblages by mapping the reads onto the consensus sequences using a minimum 50bp overlap, minimum 25 word length, and a maximum mismatch per read of 15%. Primers suitable for dPCR analyses were designed from these consensus sequences, using Primer3 software (Rozen & Skaletsky, 2000), implemented in Geneious v.9.1.7 (https://www.geneious.com). Our selection criteria for primers included an oligo size of 18 bp–22 bp, a product size of 75 bp–200 bp, a GC content of 40%–60%, a Tm of 50°C–65°C, a maximum Tm difference of 3°C. Probe selection criteria included the location between the two primers of the amplicon with no overlap with the primers, a GC content of 30%–80%, a size <30 bp, a Tm of 60°C–65°C and with 3-10°C higher than the Tm of the primers and absence of G at the 5’ end. Each probe was labeled with a distinct fluorescent dye and with BHQ1 as a quencher (Table 2). Primer specificity was verified using NCBI Primer-BLAST (Ye et al., 2012).

**Table 2.**
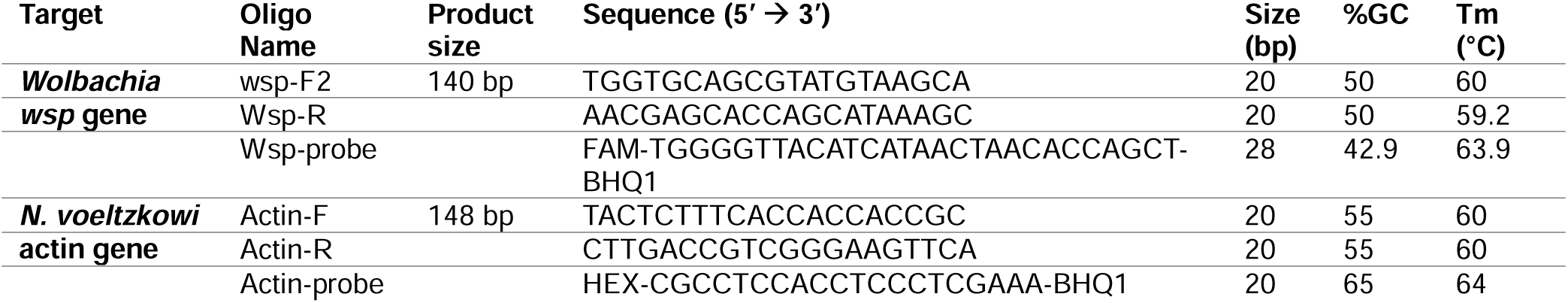
Primer and probe sequences used in the dPCR assay (Eurofins Genomics; Germany)

Quantification of the *wsp* and actin genes were performed using the QIAcuity™ One 5plex Digital PCR System (QIAGEN, Germany) according to the manufacturer’s recommendations in which 1 μl of DNA was combined with the QIAcuity Probe PCR Kit (QIAGEN, Germany) and the newly designed primers and probes (Eurofins Genomics; Germany).

In a final volume of 40 μl, each reaction mixture contained 1X probe PCR master mix, 0.8 μM forward primer, 0.8 μM reverse primer, 0.4 μM probe (for each target gene), 1 μl of template DNA and 26 μl of RNase free water. Samples were manually loaded into a 24-well 26K nanoplate, with one negative control sample well containing 1 μl RNase free water in place of the DNA. QIAGEN Priming Profile Probe PCR following specific priming for QIAcuity probe-based PCR Kits was applied to all sample types. Amplification followed the manufacturer**’**s thermal protocol: 95 °C for 2 min (initial denaturation), followed by 40 cycles of 95 °C for 15 s and 59 °C for 30 s. Partitioning, thermal cycling, and fluorescent signal processing was automatically carried out by proprietary software (QIAcuity® Software Suite Version 3.1.0.0). Imaging was conducted using the green and yellow channels with 300 and 250 ms exposure duration respectively. Positive and negative partitions were visualized and quantified automatically, and final concentrations were expressed as copies per microlitre (cp/µL).

The estimate of the *Wolbachia* titer from each individual was calculated as a ratio of the *wsp* gene to the actin gene and normalized by the original sample concentration. All control, parthenogenetic samples were in the range of 2.14-9.33 at the last time point except one sample at 19.77, which we interpret as a result of laboratory error and therefore exclude it from further analysis, see Supplementary figure 1. This sample does not change the interpretation of our results.

### Egg production in virgin sexual females

To understand if virgin sexual females are capable of laying eggs or of performing rare, spontaneous parthenogenesis (sometimes referred to as “tychoparthenogenesis”), an additional experiment was performed. Individuals of the 4^th^ larval instar of the sexual population were collected from the stock cultures and isolated each in a 3 cm Petri dish, lined with a piece of filter paper and containing a small potato sprout infested with coccids. Two days after they reached adult stage, they were sexed, and 15 females were collected for the experiment. The experiment followed the same artificial diet and coccid transfer protocol design described above, with all the females receiving an artificial diet without antibiotic. After each artificial diet period, the plugs were collected and inspected for the presence of eggs. Whenever eggs were laid in a plug, they were counted under a Nikon SMZ1000 stereo microscope and incubated as previously described above to check for development or hatching. The transfer experiment lasted for 42 (n=5) to 50 (n=10) days after which females were dissected to examine their reproductive system as previously described.

### Statistical analysis

We applied generalized linear models (GLMs) to our dataset with R version 4.4.1 (R Core Team, 2021) using a negative binomial distribution for analyzing treatment effects on egg quantity and quasibinomial distribution for egg development. Egg development was treated in the model as a binomial response using counts of eggs that either hatched or showed visual signs of development and counts of undeveloped eggs. During the treatment period, the first two time points were excluded and the total number of eggs laid by each female and the total number of those eggs that showed signs of development during that period were used to compare between the treatment groups (time points 3-8). The effects of age in days since emergence from the pupae of each female when starting the experiment and the batch in which each female was grouped were tested as predictor variables before being excluded from the models. During the observation period, the total number of eggs laid by each female were used to compare the first half of the observation period (time points 9-13) to the second half (time points 14-18).

We applied GLMs using Gaussian distribution to investigate correlation between *Wolbachia* titer estimates and age of control parthenogenetic females after the treatment and observation periods, duration of treatment for treated parthenogenetic females, and development rate of eggs. A Poisson distribution was used to analyze the effect of titer on total egg production and a quasibinomial distribution was used to analyze the effect on total egg development of control parthenogenetic females after the treatment and observation periods. Differences in *Wolbachia* titer levels between control and treated parthenogenetic females after the incomplete treatment experiment, after the treatment period, and after the observation period, as well as the difference between treated parthenogenetic females and sexual females were tested using a Welch Two Sample t-test in R version 4.4.1.

## Results

### Wolbachia titer

#### Sexually reproducing *N. voeltzkowi*

*Wolbachia wsp* gene amplification among the 13 untreated sexual females was effectively negligible, indicating a lack of infection. Sexual females had an average of 0.11±0.25 *wsp* copies per microliter, whereas the known-infected, untreated, parthenogenetic females had an average of 1,757.8 ± 782.1 with the lowest sample recording 334.0 *wsp* copies per microliter.

#### Parthenogenetic *N. voeltzkowi*

*Wolbachia* titer estimates significantly increased with the age since emergence from pupae when the control parthenogenetic females were analyzed (p= 1.6e-09). Nevertheless, titer levels in treated, parthenogenetic females were significantly lower than in control females at every time point (after the partial elimination experiment: p= 0.02, after the treatment period: p= 2.9e-14, and after the observation period: p= 0.007). In parthenogenetic females, *Wolbachia* titer estimates demonstrate that the *Wolbachia* level of treated females decreased over the course of the treatment period (time point 2 and 8) (p= 6.9e-06). While these levels were very low, they were still significantly higher than the effectively zero titer estimates of the uninfected sexual females (p= 2.8e-09), see Figure 1. After the treatment, titer levels rebounded to near control levels by the end of the observation period (time point 18) (p= 0.007), see Figure 1. We saw no effect of *Wolbachia* titer on the total quantity or quality of eggs produced by control parthenogenetic females at their last time point after the treatment period (egg production: p= 0.08, egg development: p = 0.81) or after the observation period (egg production: p= 0.12, egg development: p = 0.65), see Supplementary figure S2.

**Figure 1.**
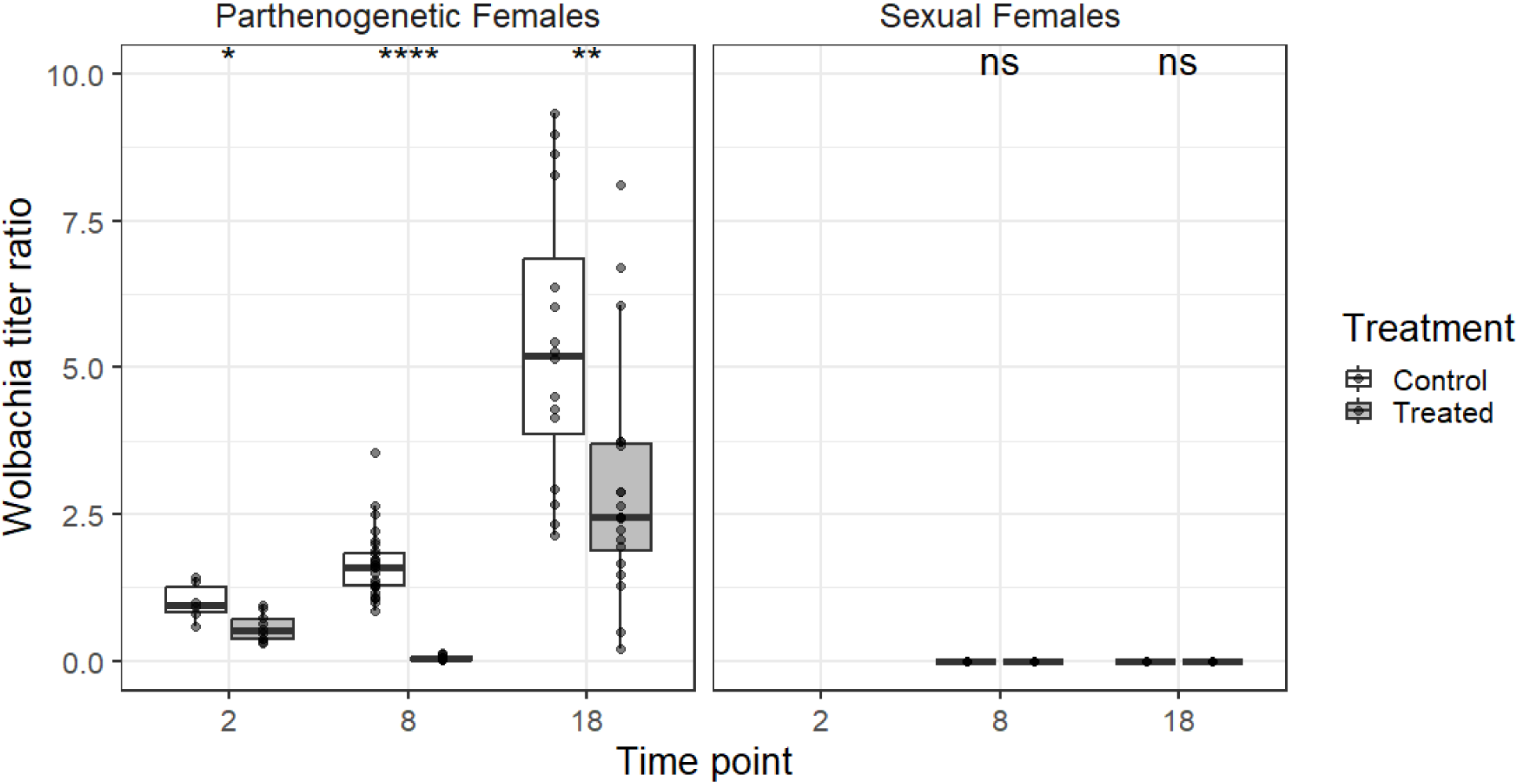
Change in *Wolbachia* titer based on the ratio of *Wolbachia wsp* gene and *N. voeltzkowi* actin gene amplification in parthenogenetic and sexual females given an antibiotic or control diet. Asterisks indicate statistical significance based on student’s t-test (ns: p > 0.05, *: p <= 0.05, **: p <= 0.01, ***: p <= 0.001, ****: p <= 0.0001).

### Antibiotic *Wolbachia* elimination and reproductive consequences

The majority of females survived until the end of the 32-day (8 time points) treatment period of the experiment (10 out of 163 females died prematurely, 6.13%, distributed across all treatment groups).

To distinguish the effect of *Wolbachia* elimination from direct toxicity effects of the antibiotic itself, we used females from the uninfected, sexual population. We did not detect any negative consequence of antibiotic treatment on egg production or development in these females. During the treatment period of the experiment, the treated, sexual females did not exhibit any significant change in total egg production or total egg development rate (egg production: p= 0.16, egg development: p=0.23, see Figure 2C) compared to the control, sexual females (see Figure 2D).

**Figure 2.**
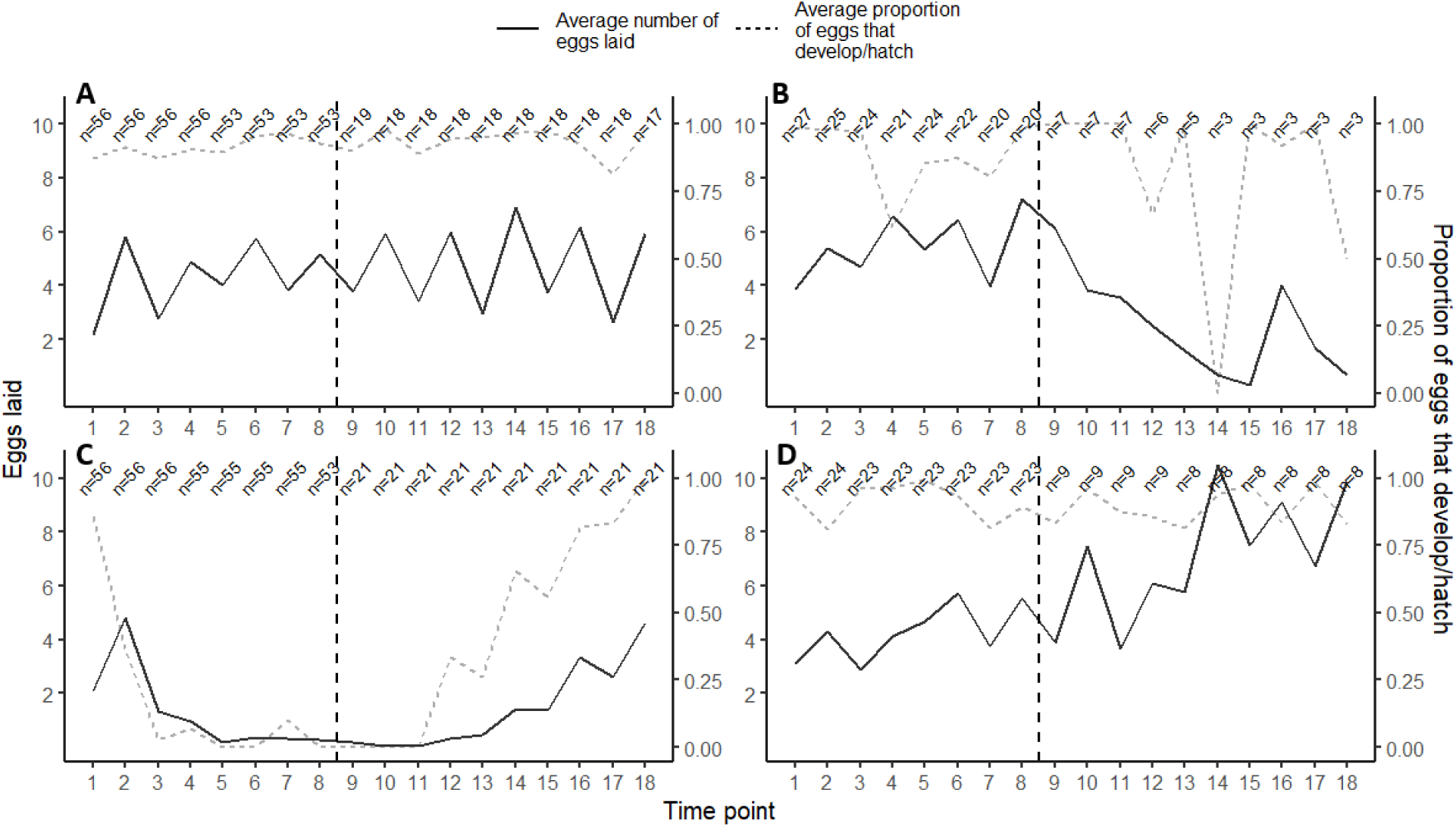
Egg quantity and quality across the treatment groups (**A**. control parthenogens, **B**. control sexuals, **C.** treated parthenogens, **D**. treated sexuals) represented by the average number of eggs produced by each female (solid line, left Y-axis) and the average proportion of those eggs that showed some sign of development (dashed line, right Y-axis), respectively. The vertical dashed line represents the end of the treatment period and start of the observation period. Sample amount corresponds to the number of females present in the experiment at that time point. Each time point is 4 days apart and eggs were laid into vial plugs for 2 days before collection.

Among the parthenogenetic females, a subgroup of the treated and untreated females was provided with a male to encourage potential mating, however, we found no significant difference in egg production or development between females that were paired with a male and those without among the treated (egg production: p= 0.75, egg development: p= 0.99) or untreated females (egg production: p=0.46, egg development: p=0.75) during the treatment period. Therefore, the two groups were pooled for further analyses.

We found that egg production and egg development rate among treated, parthenogenetic females was significantly lower (egg production: p < 2e-16, egg development: p < 2e-16, see Figure 2A) than the control, parthenogenetic females (see Figure 2B) during the treatment period. By the fourth round of treatment, the majority (56.4%) of females already stopped producing eggs and by the end of the treatment period, 73.6% of female did not produce any eggs. The remaining that did lay eggs only laid an average of 1.14 eggs, of which none hatched or showed signs of development. The age of each female at the beginning of the treatment did not have a significant effect on egg laying or development (egg production: p= 0.79, egg development: p= 0. 46). There was also no effect of the batch in which each female started the experiment on reproduction (all batches egg production: p > 0.24, all batches egg development: p > 0.43).

After 32 days, the remaining females from all treatment groups were given a control diet and followed the same feeding routine for a 38-day (10 time points) observation period. Natural mortality was still low but had slightly increased during this period (7 out of 56 females died prematurely, 12.5%, see Supplemental Table 1 for details). Over the course of the observation period, previously treated parthenogenetic individuals recovered their reproductive capabilities and the total eggs laid were significantly higher in the second half (time points 14-18) of the observation period than the first half (time points 9-13) (egg production: p < 2e-16), see Figure 2A and 2B.

Treated sexual females demonstrated an increase in average egg number towards the end of the observation period (see Figure 2C). However, the sample size of the control sexual female group became too small during the observation period, resulting in an inconsistent and likely unrepresentative average egg number.

### Wolbachia elimination effect on oogenesis

Ovary dissections were performed on a subset of females from each treatment group after the end of the treatment period (see Supplementary Table 1 for details). Parthenogenetic *N. voeltzkowi* still have eggs present in their reproductive system even after *Wolbachia* is significantly reduced by antibiotic treatment. Across all treatment groups, eggs and swollen ovariole follicles were observed in ovaries (5 out of 7 treated parthenogenetic females and 10 out of 11 control females had clear eggs in the ovaries) see example in Figure. There was no significant difference in the presence of eggs based on treatment group (p∼1). Several of the treated parthenogenetic females had well developed eggs while having not laid an egg since the 2^nd^ time point (28 days earlier), see Supplementary Table 2 for details.

### Effect of partial *Wolbachia* reduction on embryo development

Early in the experiment, when *Wolbachia* has only been slightly reduced, an average of only 36.5% (SD = 36.3%) of eggs from treated parthenogenetic females showed signs of development or hatched after the 2^nd^ antibiotic dose, whereas egg number did not yet differ from the control parthenogenetic females. In order to understand if the incomplete treatment of *Wolbachia* is affecting the development of eggs before reduction completely halts egg laying in treated parthenogenetic females, we DAPI-stained nuclei in eggs laid after the 2^nd^ antibiotic dose to see any signs of development that might be halting before the development can be seen externally. Among the 5 treated parthenogenetic females of which laid eggs were successfully retrieved, all but 1 female had eggs with developing nuclei when stained with DAPI, see **Figure 4**.

**Figure 3.**
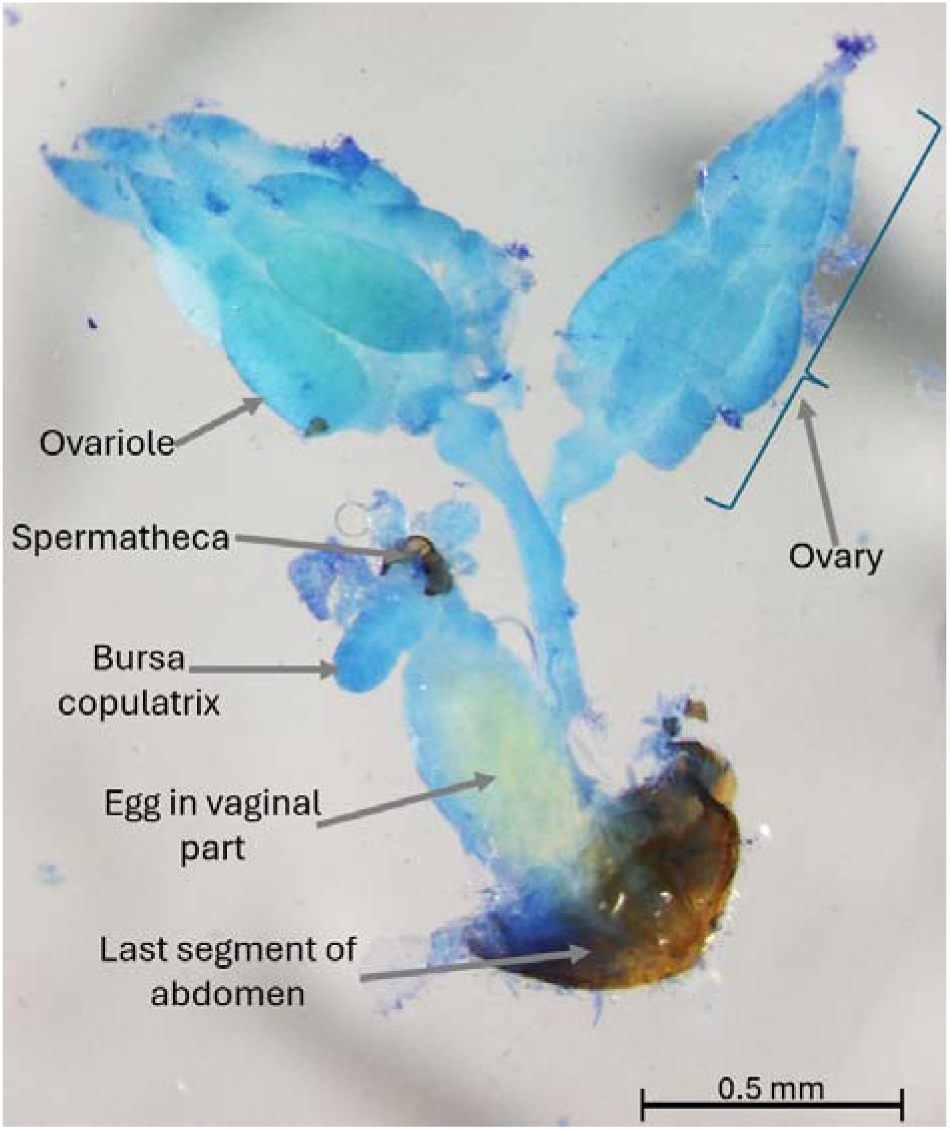
Dissection of stained reproductive system from a parthenogenetic female after antibiotic treatment. An egg is clearly present and ovarioles are well developed.

**Figure 4.**
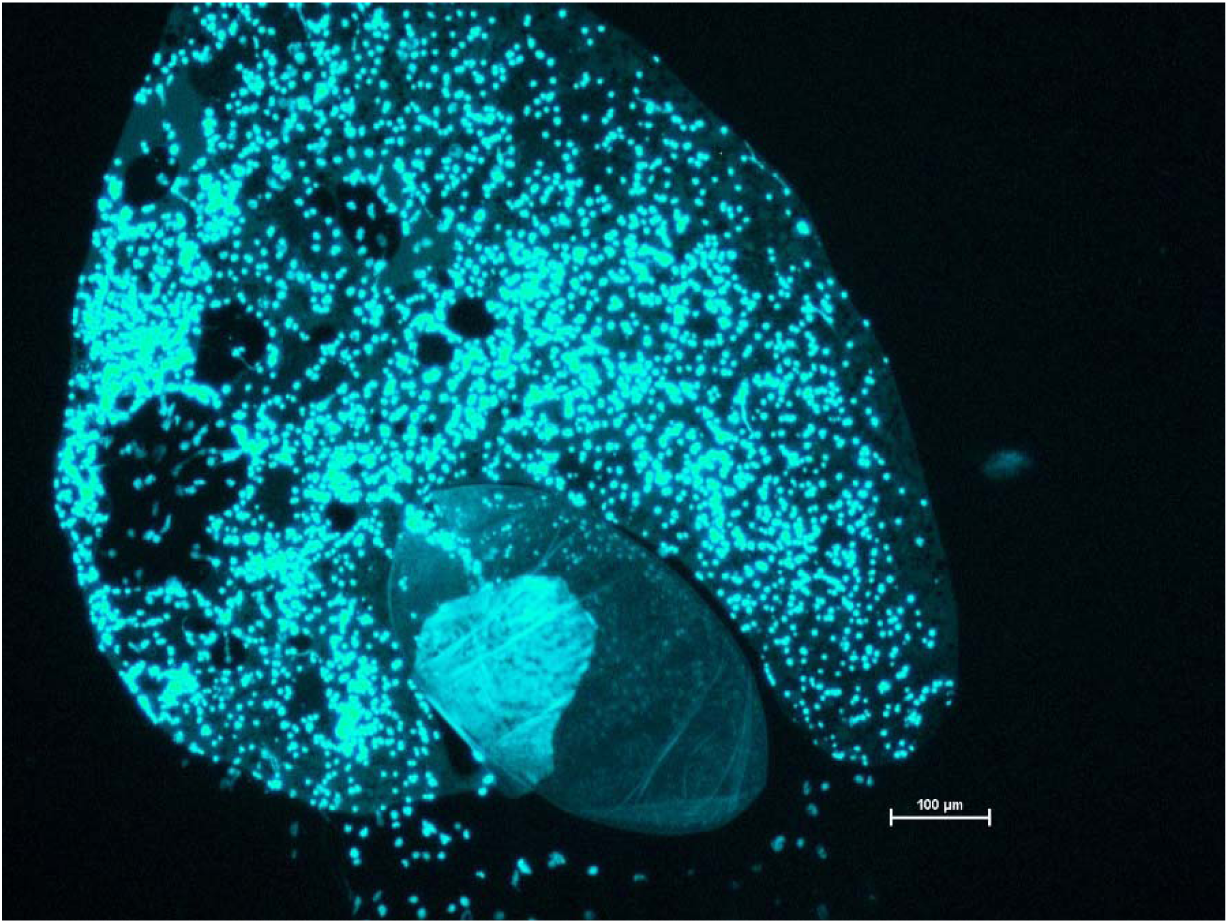
DAPI staining of a parthenogenetic female’s egg after 2 cycles of antibiotic treatment. Fluorescent nuclei are clearly visible when the egg is broken under a slide cover.

### Reproduction in virgin sexual females

We found that 5 out of 16 virgin females from the sexual population were capable of laying eggs without copulation with a male. 3 females laid only 1 egg, 1 female laid 4 eggs, and another female laid 5 eggs over the course of the 50-day of the experiment. However, none of these eggs showed any sign of development or were able to hatch. Dissection of these females showed that 8 out of 16 virgin, sexual females showed some development in ovarioles, but no female was found with a developed egg in their reproductive tract.

## Discussion

We have shown that *Wolbachia* is necessary for successful reproduction in parthenogenetic *N. voeltzkowi* females but not required in the reproduction of sexual females. As a result of treatment with Rifampicin antibiotic, we saw a significant decrease in the number of eggs oviposited by the *Wolbachia*-infected females over time, eventually resulting in the complete arrest of egg production. Paired with our results demonstrating that antibiotic treatment reduced titers of *Wolbachia*, it appears that *Wolbachia* is required in parthenogenetic *N. voeltzkowi* females in order to facilitate egg laying and development. This is further supported by the fact that we did not observe any effect of the antibiotic treatment on the uninfected, sexual *N. voeltzkowi* females; therefore, the changes in reproduction in parthenogenetic females are very unlikely a consequence of antibiotic toxicity from the treatment itself.

Early in the antibiotic treatment of the infected females, the rate of egg hatching and external signs of embryonic development in eggs laid significantly decreased compared to the control groups. Similar effects on development have been seen in *Wolbachia* elimination experiments of the parthenogenetic springtail, *Folsomia candida*, which is characterized by diplo-diploid sex determination, like *N. voeltzkowi.* However, in *Folsomia*, egg production was not affected (Pike & Kingcombe, 2009; Timmermans & Ellers, 2009). One hypothesis proposed for this result was that, if *Wolbachia* is inducing parthenogenesis via the restoration of diploidy in oocytes, the elimination of the infection would arrest this mechanism and haploid, and therefore unviable, eggs would be produced (Riparbelli et al., 2006; Timmermans & Ellers, 2009). It is unclear if this is the case with the parthenogenetic *N. voeltzkowi.* Males were provided to a subset of treated parthenogenetic females in the case that copulation may be required for oviposition or development, but we recorded no difference in any reproductive characteristic between the females paired with a mate and those without, even when instances of copulation were observed. To determine whether this reduction in egg hatching was the result of an issue with the production of the eggs prior to being oviposited or if there was a developmental issue post-oviposition, we looked for the presence of developing nuclei in recently laid eggs by parthenogenetic females in the early stage of their antibiotic treatment. As nuclei were found to be present, it appears that reduced levels of *Wolbachia* infection can affect embryonic development. A similar result was seen in a filarial nematode, *Brugia pahangi*, where reduced titers of an obligate *Wolbachia* infection via antibiotic treatment led to an increase in the amount of degenerated embryos being observed (Gunderson et al., 2020). It would appear that *Wolbachia* must be sufficiently present in order to satisfy a development checkpoint requirement to facilitate the later development of the embryo.

Our antibiotic treatment of the parthenogenetic *N. voeltzkowi* females did not fully eliminate the *Wolbachia* infection as the infection load was still higher than the uninfected, sexual females and, once the treatment stopped, the infection rebounded and reinstated reproduction. This mirrors findings from other studies of *Wolbachia* elimination with antibiotics where the *Wolbachia* infection returned to pre-treatment levels after the antibiotic administration was discontinued and their reproductive phenotypes were also reestablished (e.g. in nematodes: Gunderson et al., 2020, in springtails: Graber & Fallon, 2019; Timmermans & Ellers, 2009). How *Wolbachia* are able to resist prolonged exposure to antibiotics is not clear but may be associated with certain infected tissues being relatively impermeable to the drug, or with a protective function of specific, endosymbiont-associated tissues in the host. For example, in *Brugia pahangi* nematodes, Gunderson et al. (2020) demonstrated that dense clusters of *Wolbachia* persist in the ovaries of the host during antibiotic treatment and act as a reservoir for later proliferation and restoration of embryonic development after antibiotic treatment. Other studies have identified bacteriocyte-like cells harboring dense clusters of *Wolbachia* associated with the ovaries of the insect hosts (Hosokawa et al., 2010; Sacchi et al., 2010). These resistant clusters of *Wolbachia* could act as a backup for the infection in the event of a reduction in *Wolbachia* load.

Independently of the mechanisms preventing the complete elimination of *Wolbachia* during treatments, the persistence of low infection indicates that there must be a certain threshold that is required to facilitate the reproduction of *N. voeltzkowi*. Such thresholds appear widespread across different types of *Wolbachia*-induced reproductive manipulation (cytoplasmic incompatibility, male killing, and possibly parthenogenesis) (Bordenstein et al., 2006; Bourtzis et al., 1996; Boyle et al., 1993; Breeuwer & Werren, 1993; Fernandez Goya et al., 2026; Hurst et al., 2000; Noda et al., 2001; Rodriguero et al., 2021; Timmermans & Ellers, 2009).

The question of whether the facilitation of reproduction by *Wolbachia* in parthenogenetic *N. voeltzkowi* corresponds to the induction of parthenogenesis itself or of another aspect required for reproduction is difficult to definitively verify in non-haplo-diploid species. In haplo-diploid species, the removal of parthenogenesis-inducing *Wolbachia* results in the production of haploid gametes which are able to develop as male offspring without fertilization (Stouthamer et al., 1990; Stouthamer & Werren, 1993). When infected with *Wolbachia*, the haploid gamete genome is doubled, and in some cases feminized, by the bacteria, resulting in diploid, female offspring (Ma et al., 2015; Ma & Schwander, 2017). However, in non-haplo-diploids, after the removal of the *Wolbachia* infection needed to diploidize the gametes, they would remain haploid and would be unable to develop if unfertilized (Ma & Schwander, 2017). Reduction in egg development after the removal of a suspected parthenogenesis-inducing bacterial endosymbiont has been recorded in parthenogenetic diplo-diploid species (in weevils: Fernandez Goya et al., 2026; Rodriguero et al., 2021; Son et al., 2008, in springtails: Pike & Kingcombe, 2009; Timmermans & Ellers, 2009, and in booklice: Yusuf & Turner, 2004), but there has yet to be definitive proof that the endosymbiont induced the transition to parthenogenesis in these species. As demonstrated in our study, attempts to obtain fertilized eggs upon *Wolbachia* removal, and other studies’ attempts at the induction of parthenogenesis in sexual strains upon transfer of *Wolbachia* from the parthenogenetic ones (Kampfraath et al., 2026) have thus far been unsuccessful. In addition, there have been instances in which *Wolbachia* was found to be required for successful oogenesis in a sexual host. Thus, in the wasp, *Asobara tabida*, egg production was halted upon antibiotic treatment, without detectable effects on other traits in females or males (Dedeine et al., 2001). This indicates that the requirement of *Wolbachia* for reproduction *per se* does not mean that *Wolbachia* induces parthenogenesis in *N. voeltzkowi* or any of the above-mentioned diplo-diploid species.

Hypothetically, it is also possible for this cooccurrence of *Wolbachia* infection and parthenogenesis to occur in a system without a parthenogenesis-inducing strain of *Wolbachia*. Other mechanisms through which *Wolbachia* manipulates host reproduction to increase its own fitness, such as via bidirectional cytoplasmic incompatibility or male killing, can result in reduced host fertility and mate scarcity (Engelstädter & Telschow, 2009; Hornett et al., 2022; Hurst et al., 1999). In these cases, if a host was capable of even inefficient parthenogenesis caused by genetic factors, parthenogenesis could quickly rise in frequency in the population (Schwander et al., 2010). Such a scenario would result in a similar outcome to what is observed in *Nephus*, a cooccurrence of parthenogenesis and *Wolbachia* infection. The infection may still be necessary for successful reproduction as a leftover from its original reproductive manipulation, but this would not be direct parthenogenesis induction in its host. While bidirectional cytoplasmic incompatibility has not yet been documented in ladybugs, endosymbiont-induced male-killing has been recorded among several ladybug species (Majerus, 2006; Majerus & Hurst, 1997; Weinert et al., 2007).

While at this point, we cannot formally demonstrate that *Wolbachia* caused the transition to parthenogenesis in these populations of *N. voeltzkowi*, the cooccurrence of infection and parthenogenesis capability has thus far been consistent across *N. voeltzkowi* populations and other *Nephus* species, and we have shown that parthenogenetic reproduction is inhibited in females when *Wolbachia* is reduced. The induction of parthenogenesis by *Wolbachia* therefore seems likely. If this is a case of *Wolbachia*-induced parthenogenesis, this would not only be the first instance of parthenogenesis in Coccinellidae but also offer a novel system to understand the potential of *Wolbachia*-induced reproductive manipulation.

## Acknowledgements

We thank Jessica Ferriol for assistance with DAPI staining of eggs and microscopy imaging, Zélie Campmas for invaluable assistance in rearing and maintaining the *N. voeltzkowi* over the course of the various experiments as well as assisting with microscopy, and Jérémy Fraysse for preliminary exploration of antibiotic administration.

## Data availability

Data and scripts used in the analysis and figure generation of this article can be found at https://github.com/kjecha/Nephus_Wolbachia2026

## Notes

### Competing Interest Statement

The authors have declared no competing interest.

### Summary of Updates

Corrected a typo, and added a data availability statement

