## Supplementary tables 1-2, figure 1-2 for "*Wolbachia* facilitates the reproduction of a parthenogenetic ladybug"

### Supplementary material

##### Supplementary Table 1. Assignment of each individual into batch and treatment group based on population-associated reproductive mode (Reunion = parthenogenesis, Sète = sex). Age at start of experiment and age at death (in days since emergence from pupae) and each individual’s duration in the experiment recorded.

“End use” describes where in the analysis the specimen was used: “Wolb titer:…” – body used for *Wolbachia* titer quantification after the treatment period (time point=8) or after the observation period (time point =18), “Died” – the individual died prematurely during the experiment, days in experiment listed, “Removed due to lack of male” – the male used for the mating treatment of two female died and no other males were available as a replacement so the females were removed from the experiment, “Lost” – the individual was lost during dissection preparation, “Dissected and stained after treatment” – after 8^th^ time point, reproductive tract was dissected out, stained and photographed (see Supplementary Table 2).

| **Population** | **Treatment** | **Mate** | **Batch** | **ID #** | **Start age(days) since emergence** | **End use** | **Age(days) at death since emergence** | **Days in experiment** |
| --- | --- | --- | --- | --- | --- | --- | --- | --- |
| Reunion | Treatment | Unmated | 1 | ATU1.01 | 21 | Wolb titer: after observation | 91 | 70 |
| Reunion | Treatment | Unmated | 1 | ATU1.02 | 21 | Wolb titer: after observation | 91 | 70 |
| Reunion | Treatment | Unmated | 1 | ATU1.03 | 21 | Wolb titer: after observation | 91 | 70 |
| Reunion | Treatment | Unmated | 1 | ATU1.04 | 21 | Wolb titer: after observation | 91 | 70 |
| Reunion | Treatment | Unmated | 1 | ATU1.05 | 21 | Wolb titer: after observation | 91 | 70 |
| Reunion | Treatment | Unmated | 1 | ATU1.06 | 21 | Wolb titer: after treatment | 53 | 32 |
| Reunion | Treatment | Unmated | 1 | ATU1.07 | 21 | Wolb titer: after treatment | 53 | 32 |
| Reunion | Treatment | Unmated | 1 | ATU1.08 | 21 | Wolb titer: after treatment | 53 | 32 |
| Reunion | Treatment | Unmated | 1 | ATU1.09 | 6 | Wolb titer: after treatment | 38 | 32 |
| Reunion | Control | Unmated | 1 | ACU1.01 | 21 | Wolb titer: after observation | 91 | 70 |
| Reunion | Control | Unmated | 1 | ACU1.02 | 21 | Wolb titer: after observation | 91 | 70 |
| Reunion | Control | Unmated | 1 | ACU1.03 | 21 | Wolb titer: after observation | 91 | 70 |
| Reunion | Control | Unmated | 1 | ACU1.04 | 21 | Wolb titer: after observation | 91 | 70 |
| Reunion | Control | Unmated | 1 | ACU1.05 | 21 | Wolb titer: after treatment | 53 | 32 |
| Reunion | Control | Unmated | 1 | ACU1.06 | 21 | Wolb titer: after treatment | 53 | 32 |
| Reunion | Control | Unmated | 1 | ACU1.07 | 21 | Wolb titer: after treatment | 53 | 32 |
| Reunion | Control | Unmated | 1 | ACU1.08 | 13 | Wolb titer: after treatment | 45 | 32 |
| Reunion | Treated | Mated | 1 | ATM1.01 | 21 | Wolb titer: after observation | 91 | 70 |
| Reunion | Treated | Mated | 1 | ATM1.02 | 21 | Wolb titer: after observation | 91 | 70 |
| Reunion | Treated | Mated | 1 | ATM1.03 | 21 | Wolb titer: after observation | 91 | 70 |
| Reunion | Treated | Mated | 1 | ATM1.04 | 21 | Wolb titer: after treatment | 53 | 32 |
| Reunion | Treated | Mated | 1 | ATM1.05 | 13 | Died 26 days in | 39 | 26 |
| Reunion | Treated | Mated | 1 | ATM1.06 | 13 | Wolb titer: after treatment | 45 | 32 |
| Reunion | Control | Mated | 1 | ACM1.01 | 21 | Wolb titer: after observation | 91 | 70 |
| Reunion | Control | Mated | 1 | ACM1.02 | 21 | Wolb titer: after treatment | 53 | 32 |
| Reunion | Control | Mated | 1 | ACM1.03 | 21 | Died 14 days in | 35 | 14 |
| Reunion | Control | Mated | 1 | ACM1.04 | 21 | Wolb titer: after treatment | 53 | 32 |
| Reunion | Control | Mated | 1 | ACM1.05 | 13 | Wolb titer: after observation | 83 | 70 |
| Reunion | Control | Mated | 1 | ACM1.06 | 13 | Wolb titer: after treatment | 45 | 32 |
| Sète | Treated | Mated | 1 | STM1.01 | 22 | Died 48 days in | 70 | 48 |
| Sète | Treated | Mated | 1 | STM1.02 | 7 | Wolb titer: after treatment | 39 | 32 |
| Sète | Treated | Mated | 1 | STM1.03 | 7 | Wolb titer: after observation | 77 | 70 |
| Sète | Treated | Mated | 1 | STM1.04 | 7 | Wolb titer: after treatment | 39 | 32 |
| Sète | Treated | Mated | 1 | STM1.05 | 14 | Wolb titer: after observation | 84 | 70 |
| Sète | Treated | Mated | 1 | STM1.06 | 14 | Wolb titer: after treatment | 46 | 32 |
| Sète | Treated | Mated | 1 | STM1.07 | 14 | Wolb titer: after observation | 84 | 70 |
| Sète | Treated | Mated | 1 | STM1.08 | 14 | Wolb titer: after treatment | 46 | 32 |
| Sète | Control | Mated | 1 | SCM1.01 | 22 | Died 22 days in | 44 | 22 |
| Sète | Control | Mated | 1 | SCM1.02 | 7 | Died 22 days in | 29 | 22 |
| Sète | Control | Mated | 1 | SCM1.03 | 7 | Removed due to lack of male after 18 days | 25 | 18 |
| Sète | Control | Mated | 1 | SCM1.04 | 7 | Removed due to lack of male after 18 days | 25 | 18 |
| Sète | Control | Mated | 1 | SCM1.05 | 14 | Died 48 days in | 62 | 48 |
| Sète | Control | Mated | 1 | SCM1.06 | 14 | Wolb titer: after treatment | 46 | 32 |
| Reunion | Treatment | Unmated | 2 | ATU2.01 | 8 | Wolb titer: after observation | 78 | 70 |
| Reunion | Treatment | Unmated | 2 | ATU2.02 | 36 | Wolb titer: after observation | 106 | 70 |
| Reunion | Treatment | Unmated | 2 | ATU2.03 | 36 | Wolb titer: after observation | 106 | 70 |
| Reunion | Treatment | Unmated | 2 | ATU2.04 | 36 | Wolb titer: after observation | 106 | 70 |
| Reunion | Treatment | Unmated | 2 | ATU2.05 | 36 | Wolb titer: after observation | 106 | 70 |
| Reunion | Treatment | Unmated | 2 | ATU2.06 | 36 | Wolb titer: after observation | 106 | 70 |
| Reunion | Treatment | Unmated | 2 | ATU2.07 | 36 | Wolb titer: after treatment | 68 | 32 |
| Reunion | Treatment | Unmated | 2 | ATU2.08 | 36 | Wolb titer: after treatment | 68 | 32 |
| Reunion | Treatment | Unmated | 2 | ATU2.09 | 36 | Wolb titer: after treatment | 68 | 32 |
| Reunion | Treatment | Unmated | 2 | ATU2.10 | 36 | Wolb titer: after treatment | 68 | 32 |
| Reunion | Treatment | Unmated | 2 | ATU2.11 | 36 | Wolb titer: after treatment | 68 | 32 |
| Reunion | Treatment | Unmated | 2 | ATU2.12 | 36 | Wolb titer: after treatment | 68 | 32 |
| Reunion | Control | Unmated | 2 | ACU2.01 | 8 | Wolb titer: after observation | 78 | 70 |
| Reunion | Control | Unmated | 2 | ACU2.02 | 8 | Wolb titer: after observation | 78 | 70 |
| Reunion | Control | Unmated | 2 | ACU2.03 | 36 | Wolb titer: after observation | 106 | 70 |
| Reunion | Control | Unmated | 2 | ACU2.04 | 36 | Wolb titer: after observation | 106 | 70 |
| Reunion | Control | Unmated | 2 | ACU2.05 | 36 | Wolb titer: after observation | 106 | 70 |
| Reunion | Control | Unmated | 2 | ACU2.06 | 36 | Wolb titer: after observation | 106 | 70 |
| Reunion | Control | Unmated | 2 | ACU2.07 | 36 | Wolb titer: after treatment | 68 | 32 |
| Reunion | Control | Unmated | 2 | ACU2.08 | 36 | Wolb titer: after treatment | 68 | 32 |
| Reunion | Control | Unmated | 2 | ACU2.09 | 36 | Wolb titer: after treatment | 68 | 32 |
| Reunion | Control | Unmated | 2 | ACU2.10 | 36 | Wolb titer: after treatment | 68 | 32 |
| Reunion | Control | Unmated | 2 | ACU2.11 | 36 | Wolb titer: after treatment | 68 | 32 |
| Reunion | Control | Unmated | 2 | ACU2.12 | 36 | Wolb titer: after treatment | 68 | 32 |
| Reunion | Treated | Mated | 2 | ATM2.01 | 36 | Wolb titer: after observation | 106 | 70 |
| Reunion | Treated | Mated | 2 | ATM2.02 | 36 | Died 28 days in | 64 | 28 |
| Reunion | Treated | Mated | 2 | ATM2.03 | 36 | Wolb titer: after treatment | 68 | 32 |
| Reunion | Treated | Mated | 2 | ATM2.04 | 36 | Lost | 107 | 71 |
| Reunion | Treated | Mated | 2 | ATM2.05 | 36 | Wolb titer: after treatment | 68 | 32 |
| Reunion | Treated | Mated | 2 | ATM2.06 | 36 | Died 14 days in | 50 | 14 |
| Reunion | Treated | Mated | 2 | ATM2.07 | 36 | Wolb titer: after observation | 106 | 70 |
| Reunion | Treated | Mated | 2 | ATM2.08 | 36 | Wolb titer: after treatment | 68 | 32 |
| Reunion | Control | Mated | 2 | ACM2.01 | 36 | Died 66 days in | 102 | 66 |
| Reunion | Control | Mated | 2 | ACM2.02 | 36 | Wolb titer: after treatment | 68 | 32 |
| Reunion | Control | Mated | 2 | ACM2.03 | 36 | Wolb titer: after observation | 106 | 70 |
| Reunion | Control | Mated | 2 | ACM2.04 | 36 | Wolb titer: after treatment | 68 | 32 |
| Reunion | Control | Mated | 2 | ACM2.05 | 36 | Wolb titer: after treatment | 68 | 32 |
| Reunion | Control | Mated | 2 | ACM2.06 | 36 | Died 36 days in | 72 | 36 |
| Reunion | Control | Mated | 2 | ACM2.07 | 36 | Wolb titer: after treatment | 68 | 32 |
| Reunion | Control | Mated | 2 | ACM2.08 | 36 | Wolb titer: after observation | 106 | 70 |
| Sète | Treated | Mated | 2 | STM2.01 | 9 | Wolb titer: after treatment | 41 | 32 |
| Sète | Treated | Mated | 2 | STM2.02 | 9 | Wolb titer: after observation | 79 | 70 |
| Sète | Treated | Mated | 2 | STM2.03 | 9 | Wolb titer: after treatment | 41 | 32 |
| Sète | Treated | Mated | 2 | STM2.04 | 9 | Wolb titer: after observation | 79 | 70 |
| Sète | Treated | Mated | 2 | STM2.05 | 9 | Wolb titer: after treatment | 41 | 32 |
| Sète | Treated | Mated | 2 | STM2.06 | 9 | Died 8 days in | 17 | 8 |
| Sète | Control | Mated | 2 | SCM2.01 | 9 | Wolb titer: after treatment | 41 | 32 |
| Sète | Control | Mated | 2 | SCM2.02 | 9 | Died 50 days in | 59 | 50 |
| Sète | Control | Mated | 2 | SCM2.03 | 9 | Wolb titer: after treatment | 41 | 32 |
| Sète | Control | Mated | 2 | SCM2.04 | 9 | Died 52 days in | 61 | 52 |
| Sète | Control | Mated | 2 | SCM2.05 | 9 | Wolb titer: after treatment | 41 | 32 |
| Sète | Control | Mated | 2 | SCM2.06 | 9 | Died 42 days in | 51 | 42 |
| Sète | Control | Mated | 2 | SCM2.07 | 9 | Wolb titer: after treatment | 41 | 32 |
| Sète | Control | Mated | 2 | SCM2.08 | 9 | Wolb titer: after observation | 79 | 70 |
| Reunion | Treatment | Unmated | 3 | ATU3.01 | 10 | Wolb titer: after observation | 80 | 70 |
| Reunion | Treatment | Unmated | 3 | ATU3.02 | 10 | Wolb titer: after observation | 80 | 70 |
| Reunion | Treatment | Unmated | 3 | ATU3.03 | 10 | Wolb titer: after treatment | 42 | 32 |
| Reunion | Treatment | Unmated | 3 | ATU3.04 | 10 | Wolb titer: after treatment | 42 | 32 |
| Reunion | Control | Unmated | 3 | ACU3.01 | 10 | Wolb titer: after observation | 80 | 70 |
| Reunion | Control | Unmated | 3 | ACU3.02 | 10 | Wolb titer: after observation | 80 | 70 |
| Reunion | Control | Unmated | 3 | ACU3.03 | 10 | Wolb titer: after treatment | 42 | 32 |
| Reunion | Control | Unmated | 3 | ACU3.04 | 10 | Wolb titer: after treatment | 42 | 32 |
| Reunion | Treated | Mated | 3 | ATM3.01 | 10 | Wolb titer: after observation | 80 | 70 |
| Reunion | Treated | Mated | 3 | ATM3.02 | 10 | Wolb titer: after treatment | 42 | 32 |
| Reunion | Treated | Mated | 3 | ATM3.03 | 10 | Wolb titer: after observation | 80 | 70 |
| Reunion | Treated | Mated | 3 | ATM3.04 | 10 | Wolb titer: after treatment | 42 | 32 |
| Reunion | Control | Mated | 3 | ACM3.01 | 10 | Wolb titer: after observation | 80 | 70 |
| Reunion | Control | Mated | 3 | ACM3.02 | 10 | Wolb titer: after treatment | 42 | 32 |
| Reunion | Control | Mated | 3 | ACM3.03 | 10 | Wolb titer: after treatment | 42 | 32 |
| Reunion | Control | Mated | 3 | ACM3.04 | 10 | Died 16 days in | 26 | 16 |
| Sète | Treated | Mated | 3 | STM3.01 | 10 | Wolb titer: after observation | 80 | 70 |
| Sète | Treated | Mated | 3 | STM3.02 | 10 | Wolb titer: after treatment | 42 | 32 |
| Sète | Treated | Mated | 3 | STM3.03 | 10 | Wolb titer: after observation | 80 | 70 |
| Sète | Treated | Mated | 3 | STM3.04 | 10 | Wolb titer: after treatment | 42 | 32 |
| Sète | Treated | Mated | 3 | STM3.05 | 10 | Wolb titer: after observation | 80 | 70 |
| Sète | Treated | Mated | 3 | STM3.06 | 10 | Wolb titer: after treatment | 42 | 32 |
| Sète | Control | Mated | 3 | SCM3.01 | 10 | Wolb titer: after observation | 80 | 70 |
| Sète | Control | Mated | 3 | SCM3.02 | 10 | Wolb titer: after treatment | 42 | 32 |
| Sète | Control | Mated | 3 | SCM3.03 | 10 | Wolb titer: after observation | 80 | 70 |
| Sète | Control | Mated | 3 | SCM3.05 | 10 | Removed due to lack of male after 5 days | 15 | 5 |
| Sète | Control | Mated | 3 | SCM3.06 | 10 | Removed due to lack of male after 5 days | 15 | 5 |
| Reunion | Treatment | Unmated | 4 | ATU4.01 | 10 | Wolb titer: after treatment | 42 | 32 |
| Reunion | Treatment | Unmated | 4 | ATU4.02 | 10 | Dissected and stained after treatment | 44 | 34 |
| Reunion | Treatment | Unmated | 4 | ATU4.03 | 10 | Dissected and stained after treatment | 45 | 35 |
| Reunion | Control | Unmated | 4 | ACU4.01 | 10 | Wolb titer: after treatment | 42 | 32 |
| Reunion | Control | Unmated | 4 | ACU4.02 | 10 | Wolb titer: after treatment | 42 | 32 |
| Reunion | Control | Unmated | 4 | ACU4.03 | 10 | Dissected and stained after treatment | 44 | 34 |
| Reunion | Control | Unmated | 4 | ACU4.04 | 10 | Dissected and stained after treatment | 45 | 35 |
| Reunion | Treated | Mated | 4 | ATM4.01 | 10 | Wolb titer: after treatment | 42 | 32 |
| Reunion | Treated | Mated | 4 | ATM4.02 | 10 | Dissected and stained after treatment | 44 | 34 |
| Reunion | Treated | Mated | 4 | ATM4.03 | 10 | Wolb titer: after treatment | 42 | 32 |
| Reunion | Treated | Mated | 4 | ATM4.04 | 10 | Dissected and stained after treatment | 45 | 35 |
| Reunion | Treated | Mated | 4 | ATM4.05 | 10 | Wolb titer: after treatment | 42 | 32 |
| Reunion | Treated | Mated | 4 | ATM4.06 | 10 | Dissected and stained after treatment | 45 | 35 |
| Reunion | Control | Mated | 4 | ACM4.01 | 10 | Wolb titer: after treatment | 42 | 32 |
| Reunion | Control | Mated | 4 | ACM4.02 | 10 | Dissected and stained after treatment | 44 | 34 |
| Reunion | Control | Mated | 4 | ACM4.03 | 10 | Wolb titer: after treatment | 42 | 32 |
| Reunion | Control | Mated | 4 | ACM4.04 | 10 | Dissected and stained after treatment | 45 | 35 |
| Reunion | Control | Mated | 4 | ACM4.05 | 10 | Died 26 days in | 36 | 26 |
| Reunion | Control | Mated | 4 | ACM4.06 | 10 | Wolb titer: after treatment | 42 | 32 |
| Sète | Treated | Mated | 4 | STM4.01 | 11 | Wolb titer: after treatment | 43 | 32 |
| Sète | Treated | Mated | 4 | STM4.02 | 11 | Dissected and stained after treatment | 45 | 34 |
| Sète | Treated | Mated | 4 | STM4.03 | 11 | Wolb titer: after treatment | 43 | 32 |
| Sète | Treated | Mated | 4 | STM4.04 | 11 | Dissected and stained after treatment | 46 | 35 |
| Sète | Control | Mated | 4 | SCM4.01 | 11 | Wolb titer: after treatment | 43 | 32 |
| Sète | Control | Mated | 4 | SCM4.02 | 11 | Dissected and stained after treatment | 45 | 34 |
| Sète | Control | Mated | 4 | SCM4.03 | 11 | Wolb titer: after treatment | 43 | 32 |
| Sète | Control | Mated | 4 | SCM4.04 | 11 | Dissected and stained after treatment | 46 | 35 |
| Reunion | Treatment | Unmated | 5 | ATU5.01 | 11 | Wolb titer: after treatment | 43 | 32 |
| Reunion | Treatment | Unmated | 5 | ATU5.02 | 11 | Dissected and stained after treatment | 44 | 33 |
| Reunion | Control | Unmated | 5 | ACU5.01 | 11 | Wolb titer: after treatment | 43 | 32 |
| Reunion | Control | Unmated | 5 | ACU5.02 | 11 | Dissected and stained after treatment | 44 | 33 |
| Reunion | Treated | Mated | 5 | ATM5.01 | 11 | Wolb titer: after treatment | 43 | 32 |
| Reunion | Treated | Mated | 5 | ATM5.02 | 11 | Dissected and stained after treatment | 44 | 33 |
| Reunion | Control | Mated | 5 | ACM5.01 | 11 | Wolb titer: after treatment | 43 | 32 |
| Reunion | Control | Mated | 5 | ACM5.02 | 11 | Dissected and stained after treatment | 44 | 33 |
| Sète | Control | Mated | 5 | SCM5.01 | 12 | Wolb titer: after treatment | 44 | 32 |
| Sète | Control | Mated | 5 | SCM5.02 | 12 | Dissected and stained after treatment | 45 | 33 |
| Sète | Control | Mated | 5 | SCM5.03 | 12 | Wolb titer: after treatment | 44 | 32 |
| Sète | Control | Mated | 5 | SCM5.04 | 12 | Died 8 days in | 20 | 8 |

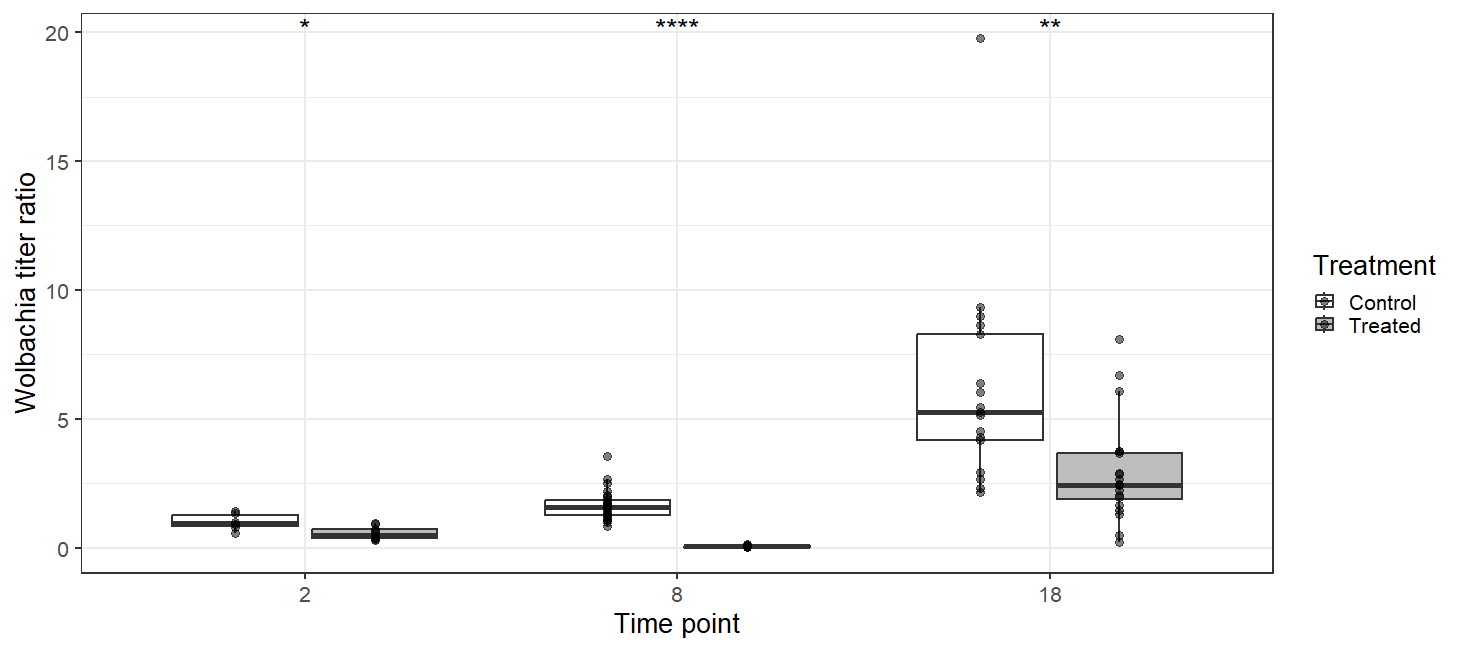

##### Supplementary Figure 1. *Wolbachia* titer including outlier data point ( time point = 18, control treatment)

Change in *Wolbachia* titer based on the ratio of *Wolbachia* *wsp* gene and *N. voeltzkowi* actin gene amplification in parthenogenetic and sexual females given an antibiotic or control diet. Asterisks indicate statistical significance based on student’s t-test (ns: p > 0.05, *: p <= 0.05, **: p <= 0.01, ***: p <= 0.001, ****: p <= 0.0001). No change in significance when outlier is included.

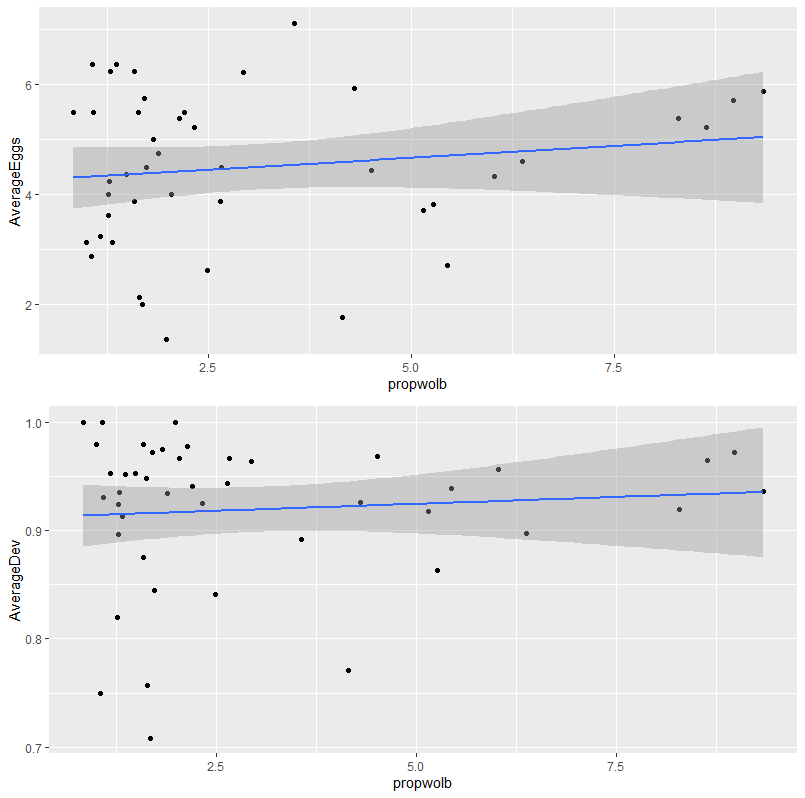

##### Supplementary Figure 2. No effect of *Wolbachia* titer on egg production or development.

“AverageEggs” and “AverageDev” are the average number of eggs produced and proportion of those eggs that developed, respectively, by each female over the course of the experiment, only considering days active in the experiment. “Propwolb” is the calculated as a ratio of the *Wolbachia* *wsp* gene to the *Nephus* actin gene and normalized by the original sample concentration.

##### Supplementary Table 2. Ovary dissection results at end of treatment period (time point = 8).

ID’s denote treatment group: A/S = asex/sex, C/T= control/treated, M/U= mated/unmated. Presence of mature eggs and developed ovarioles recorded. Time points of last egg laid also recorded. Treated, parthenogenetic females in bold.

| **ID** | **Eggs?** | **Developed ovarioles?** | **Last egg laid time point (dissected after TP=8)** | **picture** | **notes** |
| --- | --- | --- | --- | --- | --- |
| ACM4.02 | Yes | No | 8 | 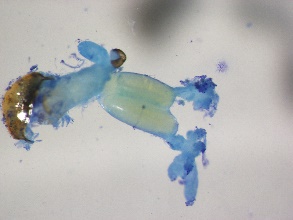 | 2 eggs stuck in oviduct, shriveled ovaries |
| ACM4.04 | Yes | No | 8 | 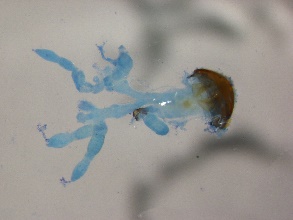 | broke ovary while dissecting but saw an egg |
| ACM5.02 | Yes | Yes | 6 | 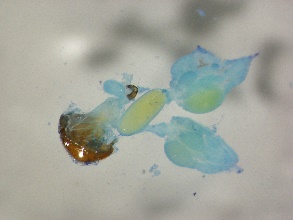 |  |
| ACU4.03 | Yes | Yes | 8 | 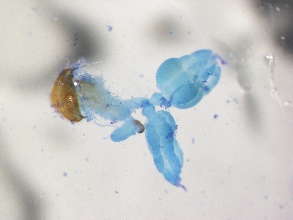 |  |
| ACU4.04 | Yes | Yes | 8 | 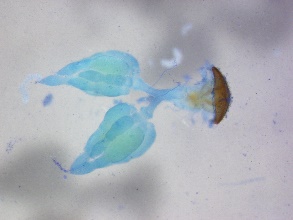 |  |
| ACU5.02 | Yes | No | 6 | 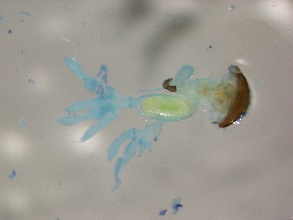 |  |
| **ATM4.02** | Yes | Yes | 8 | 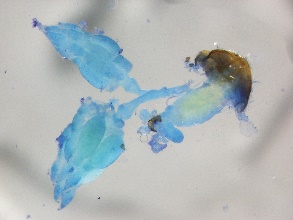 |  |
| **ATM4.04** | Yes | Yes | 3 | 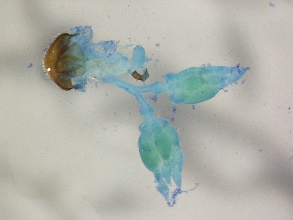 | squeezed egg out while dissecting |
| **ATM4.06** | Yes | Yes | 4 | 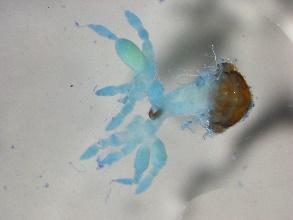 |  |
| **ATM5.02** | No | No | 6 | 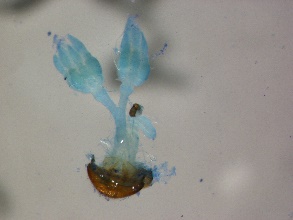 |  |
| **ATU4.02** | Yes | Yes | 3 | 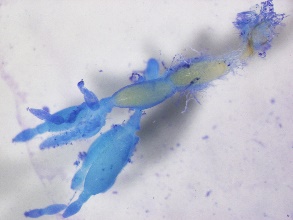 | 2 eggs in oviduct |
| **ATU4.03** | Yes | Yes | 2 | 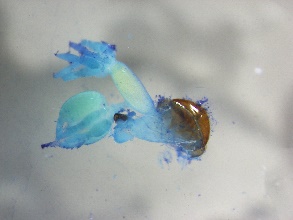 |  |
| **ATU5.02** | No? | Yes | 6 | 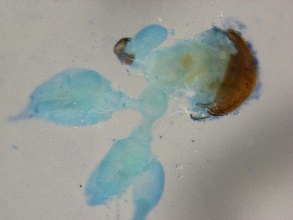 | yellow ball in oviduct but not oval shaped like egg |
| SCM4.02 | Yes | Yes | 8 | 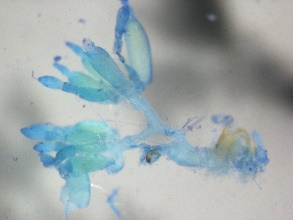 |  |
| SCM4.04 | Yes | Yes | 8 | 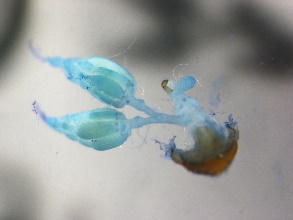 |  |
| SCM5.02 | Yes | Yes | 8 | 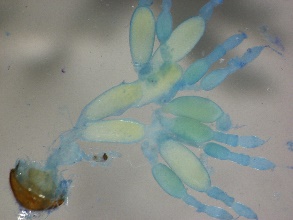 |  |
| STM4.02 | Yes | Yes | 8 | 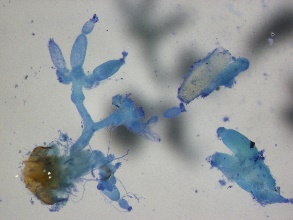 | broke ovary while dissecting but saw an egg |
| STM4.04 | No | No | 2 | 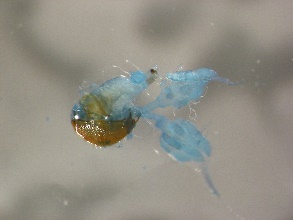 | no eggs but hasn't laid egg recently |
